# Quantitative comparison of single-cell sequencing methods using hippocampal neurons

**DOI:** 10.1101/004366

**Authors:** Luwen Ning, Guan Wang, Zhoufang Li, Wen Hu, Qingming Hou, Yin Tong, Meng Zhang, Li Qin, Xiaoping Chen, Hengye Man, Pinghua Liu, Jiankui He

## Abstract

Single-cell genomic analysis has grown rapidly in recent years and will find widespread applications in various fields of biology, including cancer biology, development, immunology, pre-implantation genetic diagnosis, and neurobiology. In this study, we amplified genomic DNA from individual hippocampal neurons using one of three single-cell DNA amplification methods (multiple annealing and looping-based amplification cycles (MALBAC), multiple displacement amplification (MDA), and GenomePlex whole genome amplification (WGA4)). We then systematically evaluated the genome coverage, GC-bias, reproducibility, and copy number variations among individual neurons. Our results showed that single-cell genome sequencing results obtained from the MALBAC and WGA4 methods are highly reproducible and have a high success rate. Chromosome-level and subchromosomal-level copy number variations among individual neurons can be detected.

## INTRODUCTION

Because of somatic mutations, the genomes of two cells are unlikely to be identical(Frumkin et al. 2005). Biologists have long been interested in analyzing genomic diversity among individual cells. Unlike traditional DNA sequencing, in which genetic materials are isolated from a population of cells, amplification of the genetic material from a single cell by a high-fidelity and low-bias method is key to the success of single-cell genome sequencing studies. Over the years, several single-cell whole genome amplification methods were reported. The first method is multiple displacement amplification (MDA). MDA is a non-PCR-based DNA amplification technique, which uses a high fidelity enzyme, preferentially Φ29 DNA polymerase, to amplify the target genome(Dean et al. 2002; Spits et al. 2006a),^9^. The MDA method has been applied to study genetic diversity among individual cells in solid tumors(Xu et al. 2012). GenomePlex whole genome amplification (WGA4) is another single-cell whole genome amplification method, which is based on the PCR amplification of randomly fragmented genomic DNAs using universal oligonucleotides as primers. Recently, WGA4 was applied to analyze cancer cell copy number variation (CNV)(Navin et al. 2011). The WGA4 method has also been used to study genomic diversity among neurons^10^. Last year, Zong *et al*. described a third single-cell genome amplification method, the multiple annealing and looping-based amplification cycles (MALBAC) method(Zong et al. 2012). The MALBAC method was able to achieve up to 93% coverage of the human genome at a 25× mean sequencing depth^6^.

Single-cell whole genome sequencing has been applied to study cancer biology, cell development, neurobiology, and pre-implantation genetic diagnosis(Ramsköld et al. 2012; Wang et al. 2012; Yan et al. 2013)^10^,^11^. In the human brain, the ≈85 billion individual neurons(Williams and Herrup 1988; Azevedo et al. 2009) show remarkable diversity in their maturation, morphology, electrophysiological properties, and interneuronal connectivity. Somatic variations of the genome and epigenome, including chromosome instability, aneuploidy (rarely polyploidy), mosaic subchromosomal rearrangements, and changes in epigenetic modifications, all contribute to the creation of neuronal diversity(Muotri and Gage 2006). Thus, neurons are a suitable system to study single-cell genome diversity. In this study, after the quality of single-neuron genome sequencing was confirmed by comparison to the results of traditional sequencing studies (using genomic DNA from ≈2 million neurons of the same rat, abbreviated as bulk cells in this study),we quantitatively analyzed 11 neurons using the MDA, WGA4, and MALBAC techniques with an emphasis on the following questions: 1) At the same sequencing depth, which of the three methods gives the best genome coverage? 2) Is there amplification bias among different genomic regions, and can the bias issue be addressed? 3) How reproducible are these three whole genome amplification methods? 4) What are the major advantages for each of the three single-cell whole genome amplification methods? Our results demonstrated that single-cell genome sequencing results using either the MALBAC or WGA4 method are highly reproducible and have a high success rate. Chromosome-level and subchromosomal-level CNVs among individual neurons can be detected.

## RESULTS

### Experiment Design

The general strategies that were used in sample preparation, DNA sequencing, and data analysis are summarized in Fig. 1. Hippocampal neurons were prepared from individual E18 rat embryos and cultured in neurobasal medium as described previously(Zhang et al. 2009; Hou et al. 2011). Nuclei of individual hippocampal neurons were collected using a glass micropipette (Fig.1b,c)(Campbell et al. 2006; Nawy 2014) and transferred directly to 200-µL PCR tubes. The single neuron nucleus was then subjected to whole genome amplification using one of three methods (2 nuclei by MDA, 5 nuclei by MALBAC, and 4 nuclei by WGA4). At the same time, genomic DNA was also isolated from 2 million cultured neurons (bulk cells) derived from the same embryos. Sequencing libraries were constructed following the Illumina standard protocol and sequenced by an Illumina HiSeq 2000. On average, there were 34 million clean reads per sample (**Supplementary Table S1**). The data were mapped to the rat reference genome (rnt5) using Bowtie2(Benjamini and Speed 2012). Using the sequencing results from bulk cells as the benchmarks, single-cell genome sequencing results were analyzed for genome coverage, GC-biases, reproducibility, and CNV(Ning et al. 2014).

**Figure 1.**
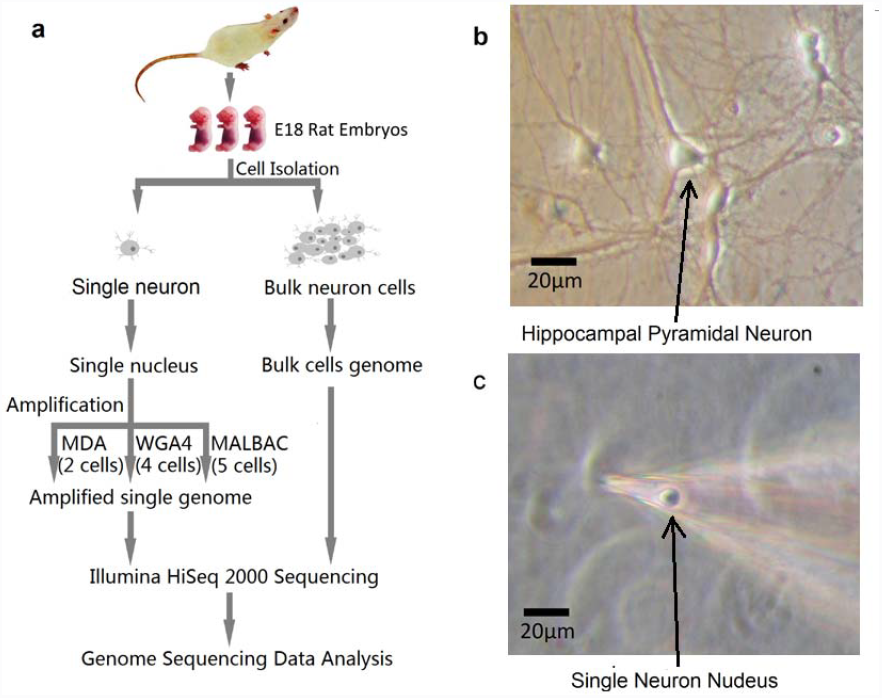
Experimental design. (**a**) Flow chart of bulk-neuron and single-neuron genomic isolation, DNA sequencing sample preparation, sequencing, and data processing. (**b-c**) Culture and isolation of the nucleus from individual neurons. The nucleus was extracted directly from cultured neurons through micromanipulation using a micro glass pipette on an electrophysiological recording system. A picture of the isolated nucleus in the micropipette is shown (**c**).

### Genome coverage efficiency

Genome coverage is the percentage of the genome that is covered by at least a single base after aligning the sequence data to the reference genome. Because genome amplification is a crucial step in single-cell genome sequencing studies, some of the major concerns include whether the genome amplification is uniform and whether certain genomic regions may be lost during genome amplification. Current single-cell whole genome sequencing studies usually have a low sequencing depth (< 1×)(Navin et al. 2011; Evrony et al. 2012; McConnell et al. 2013). Therefore, it is important to evaluate the performance of single-cell amplification/sequencing techniques at low sequencing depth (< 1×). As shown in Fig 2, genome sequencing using DNA from bulk cells achieved a coverage of ≈0.5 at a sequencing depth of 1×. Genome coverage also improves with the increasing of sequencing depth. At the same level of sequencing depth (< 1×), the results from bulk cells have significantly better coverage than any of the single-cell amplification/sequencing methods. For all three single-cell genome amplification methods, the genome coverage also improves when the sequencing depth increases. Among the three single-cell amplification methods, MALBAC has the best performance in terms of genome coverage, which is consistent with the observation of a previous report(Zong et al. 2012). At a sequencing depth of 1×, the genome coverage from MALBAC can be as good as 0.3 (Fig. 2).

**Figure 2.**
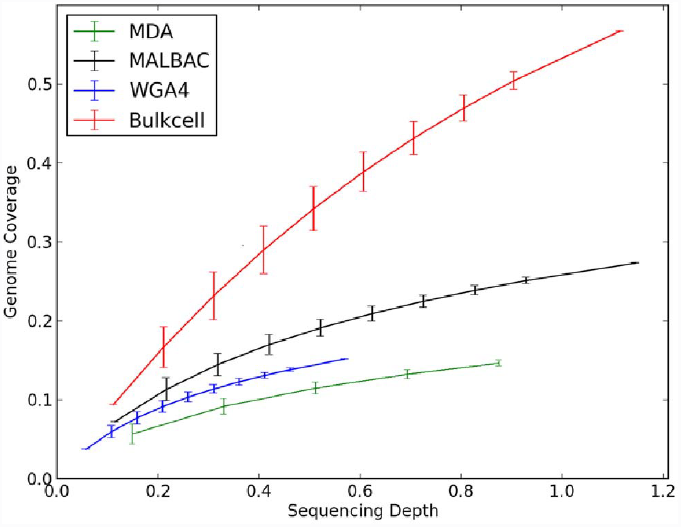
Genome coverage increases with sequencing depth. For each single-cell amplification/sequencing method evaluated, a representative cell having the best coverage was used (see **Supplementary Fig. S1** for the information of other individual cells used in this study).The error bars indicate the standard deviation of 10 replicates of sampling when the coverage was calculated.

### GC-bias

GC-bias is a parameter that is used to quantitatively evaluate whether there is a correlation between the observed coverage of a specific genomic region and its GC content. The presence of GC-bias can complicate data analysis, including copy number estimation. In single-cell whole genome sequencing studies, GC-biases may result from either the whole genome amplification step or the sequencing step. In our study, before analyzing individual cells, the sequencing results of DNA from bulk cells were used as the benchmark to quantitatively evaluate the sequencing platform because this sample was sequenced without the need for genome amplification. The average GC content of bulk-cell genomic DNA sequencing data is 41.4%, which is 0.5% less than the GC content of the reference genome (41.9%) (Fig. 3a). Previous studies suggested that the GC-rich regions are prone to low coverage with the Illumina HiSeq 2000 platform(Venkatraman and Olshen 2007). Our results indicated that there is a very low level of GC-bias when sequencing the rat genome with the Illumina HiSeq 2000 platform.

**Figure 3.**
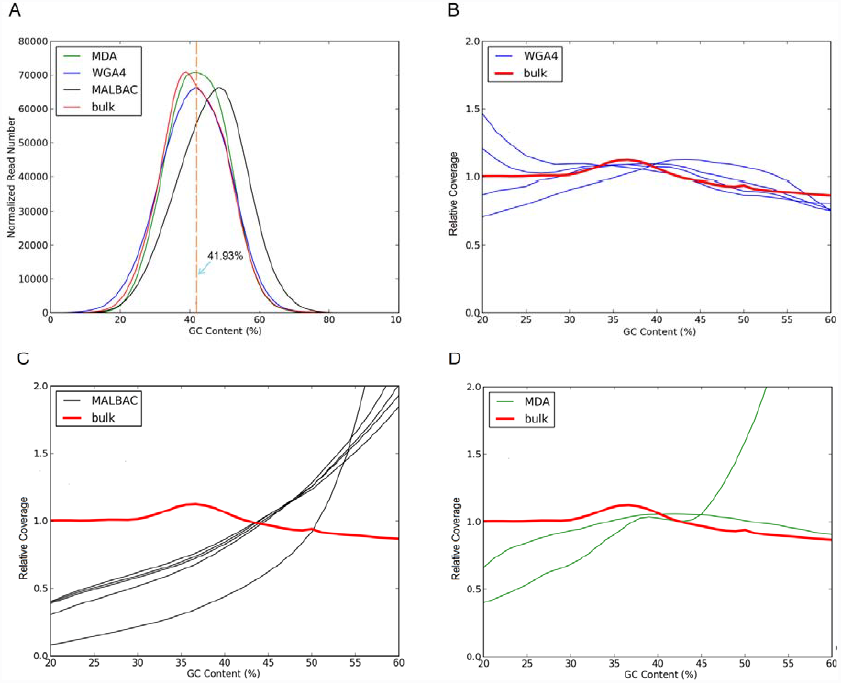
GC content analysis. (**a**) The GC-composition of four samples (bulk-cell sample and DNA amplified by three different single-cell amplification methods). The y-axis is the frequency of normalized reads of with different GC contents. The total reads of each sample is normalized to 10 million. The average GC content of the rat reference genome is 41.9% (dashed line). The average GC content of the MDA, WGA4, MALBAC, and unamplified samples are 43.4%, 41.6%, 46.6%, and 41.4%, respectively. (**b-d**) The GC-bias plot. The relative coverage (y-axis) represents the ratio between the coverage of a sample and the coverage predicted by the reference genome. A relative coverage of 1 indicates no bias. A relative coverage above 1 or below 1 indicates higher or lower coverage than that expected, respectively. The results from the bulk-cell sample (red line) are plotted as a benchmark.

Because of the low level of GC-bias at the sequencing step on the Illumina HiSeq 2000 platform, we mainly focused on the GC-bias during the whole genome amplification step. The average GC content (41.6%) of the DNA samples amplified from a single cell using the WGA4 method is very close to that of the reference genome (41.9%), with a difference of ≈0.3% (Fig.3a). The average GC content (43.4%) of DNA prepared by the MDA method is ≈1.5% higher than that of reference genome. However, the average GC content (46.6%) of DNA prepared by the MALBAC method is ≈4.7% greater than that of reference genome, indicating some degree of preference for GC-rich regions during the amplification process. After examining the overall GC-bias, we then analyzed the relationship between the observed reads and the GC content of individual genomic regions (Fig. 3b–d). The majority of rat genomic sequences (75%) have a GC content of 20–60%. Using the sequencing results from bulk cells, the plot of the relative coverage at various genomic regions versus the GC content (20–60%) is almost uniform with a very small bias toward regions having ≈35–40% GC. This data provide further evidence that the Illumina HiSeq 2000 platform has a very low level of GC-bias for rat genome sequencing studies. For single-cell genomes amplified using the WGA4 method, regions with ≈40–60% GC content have low GC-bias, and the cell-to-cell results are also highly reproducible. However, a significant level of cell-to-cell variation exists at the low-GC-content regions (< 30% GC content, Fig. 3b). MALBAC has a clear preference for high-GC-content regions relative to the low-GC-content regions (Fig. 3c). Also, among the five cells amplified using MALBAC, results from four of the five cells follow a very similar pattern. Genomic regions with a high-GC-content tended to be over-amplified when the MALBAC method was used (Fig. 3c and Supplementary Fig. S5). High-GC-content fragments may form loop structures more efficiently, which explains the systematic bias of MALBAC toward high-GC-content regions. A similar GC-bias was also noticed when MALBAC was applied to characterize human cancer cell lines(Zong et al. 2012). The two MDA cells had completely different patterns when GC-bias was analyzed, where one cell displayed a strong GC-bias while the other cell did not. Such a high level of cell-to-cell variation was also observed in studies from other groups when MDA was used as the amplification method(Rodrigue et al. 2009; Yilmaz et al. 2010; Ellegaard et al. 2013) (**Supplementary Fig. S6**). The high cell-to-cell variation may be related to the mechanism of the MDA method; depending on where the hexamer primers bind, the MDA process is random in nature(Spits et al. 2006b). Because of the high level of cell-to-cell variation in our study and the results reported in the literature(Rodrigue et al. 2009; Yilmaz et al. 2010; Ellegaard et al. 2013), we analyzed only two neurons by the MDA method. As a result, we did not analyze CNVs in these two neurons. In addition, CNV information that was reported in MDA-based single-cell studies might need to be used with caution.

In recent years, single-cell sequencing has been applied to sequence the genome of uncultured bacteria and archaea(Chitsaz et al. 2011; Yoon et al. 2011; Stepanauskas 2012). The GC content of microorganism genomes ranges from 16% to 75% in organisms such as *Candidatus Zinderia insecticola* (13.5%), *Plasmodium falciparum* (mean 19%), *Escherichia coli* (51%), and *Rhodobacter sphaeroides* (69%)(Guo et al. 2009). High-GC-content genomes are traditionally difficult to sequence. MALBAC displays a preference for high-GC-content regions, and it may have advantages in sequencing high-GC-content organisms. WGA4 may be suitable for sequencing normal and low-GC-content organisms.

### Reproducibility

Because each cell has only a tiny amount of DNA that is typically measured in picograms, the success of each experiment highly depends on the quality of the sample, the skills of operators, and potential contamination in the lab. To extract useful information from single-cell genomic studies, one of the key issues to be addressed is to distinguish between true biological cell-to-cell variation and non-specific experimental noise or errors. Thus, the next question addressed in our study was that of the cell-to-cell genome amplification reproducibility. To evaluate the reproducibility of the three whole genome amplification methods (MDA, WGA4,and MALBAC), the rat reference genome was partitioned into 500 kb-sized bins, yielding 5,797 bins in total. After calculating the reads mapped to each bin, we then compared the number of normalized reads of each bin between two cells (Fig. 4a–c). Our data show that amplifications by both MALBAC and WGA4 are highly reproducible, with the correlation coefficient > 0.9 for the MALBAC method and close to 0.9 for the WGA4 method. Such high correlation coefficients suggest that single-cell genome amplification by either the MALBAC or WGA4 method is highly reproducible and that the information extracted from single-cell genomic studies can be used to analyze cell-to-cell genomic diversity. We also calculated the correlation coefficient matrix between any two cells among the 11 single cells that were analyzed in our studies and their correlation with the sequencing results from bulk cells. Hierarchical clustering of the correlation coefficient matrix shows that single cells from the same method are clustered together (Fig. 4d). However, cells that were amplified using different methods display poor correlation, indicating that each method has its own built-in pattern of biases. Thus, single-cell genomes that are amplified using different methods may not be suitable for comparative studies. Because of their high reproducibility, very useful information might be extracted from single-cell genomic studies using either the WGA4 or MALBAC method once proper GC-bias and other built-in patterns of bias are considered.

**Figure 4.**
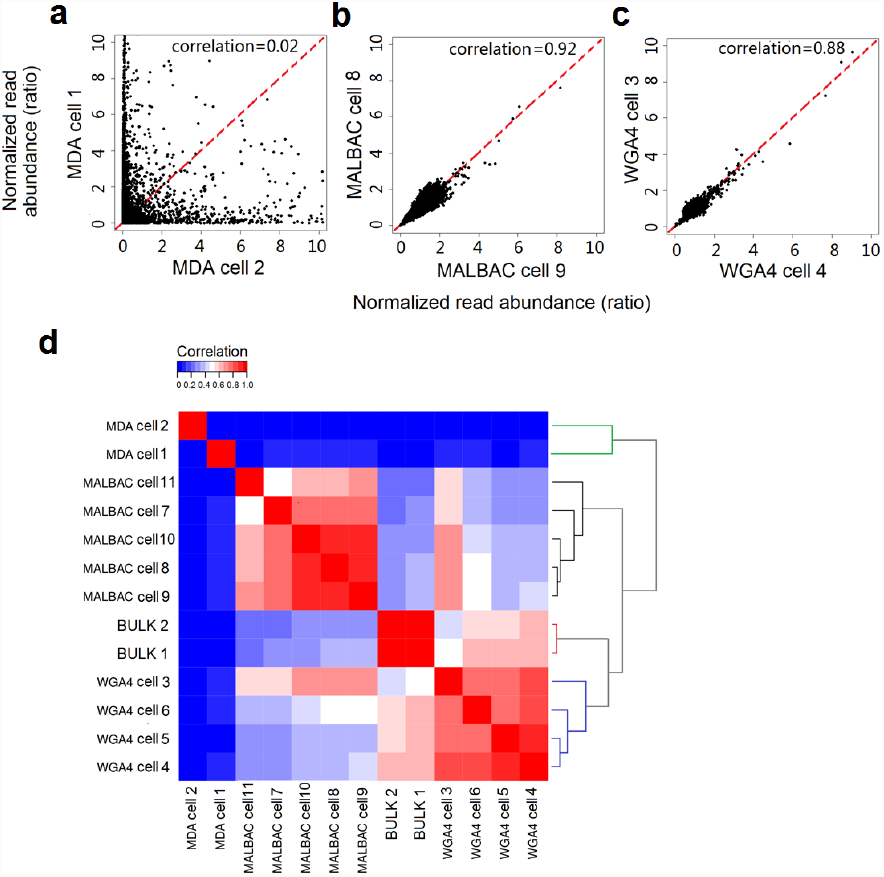
Reproducibility of three different whole genome amplification methods. (**a-c**) For each bin that is 500 kb in size, the normalized read abundance of one cell is the x-axis value, and the normalized read abundance of another cell is the y-axis value. The combination of these values will give one dot on the plot for a particular bin. There are 5,797 bins in total. The 5,797 dots are plotted to show the reproducibility of the three single-cell genome amplification methods. A narrow distribution of dots along the y = x (red line) indicates good correlation between the two cells. See **Supplementary Fig. S2** for all cells used in this study and **Supplementary Fig. S3** for results when a 200-kb bin size was used. (**d**) Hierarchical clustering is performed on the correlation of each of the single cells used in this study and the bulk-cell sample.

### Genome coverage uniformity

The genome coverage uniformity represents the evenness of sequence read distribution over the entire genome. For samples amplified using the MDA method, the sequence reads tend to concentrate at some genomic regions, and most genomic regions have relatively few reads. As evidenced by the spike observed in Fig 5a, reads from MDA samples have a very high level of bin-to-bin variation. Among the three single-cell genome amplification methods examined in this study, samples amplified using the WGA4 method showed the smallest bin-to-bin variation in read abundance (Fig. 5a), which is consistent with the GC-bias analysis shown in Fig. 3b.

**Figure 5.**
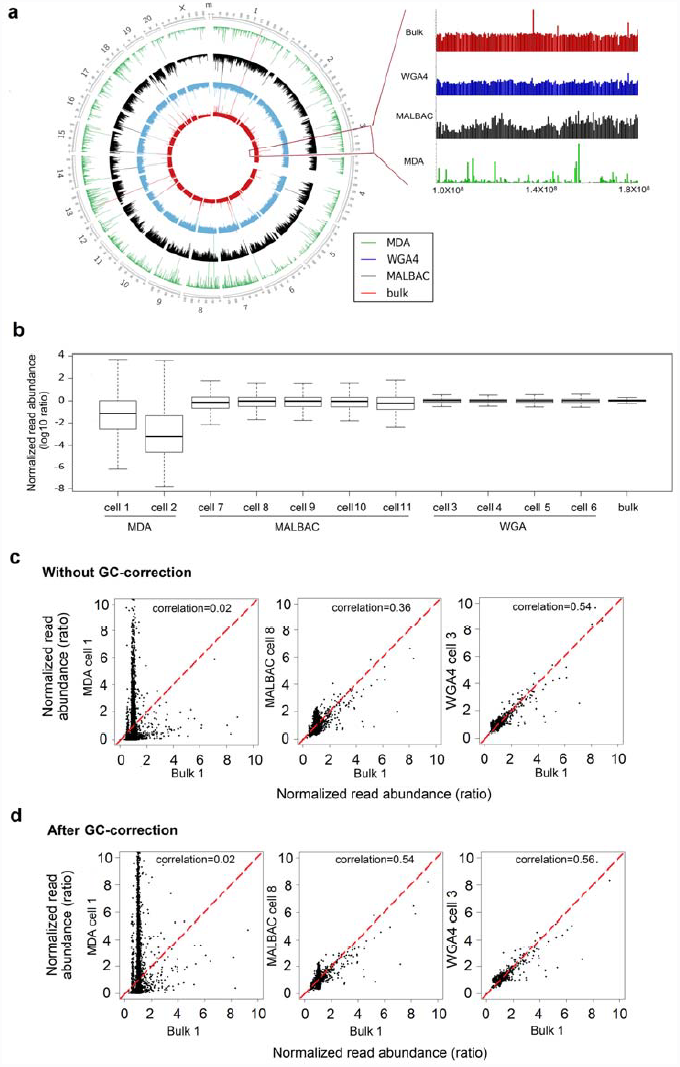
Read abundance. (**a**) Read abundance distribution across the genome from three single-cell amplification methods and a bulk-cell sample (left) and a magnified plot of part of chromosome 3 (right). The five circles from outside to inside are the chromosome position index, read abundance of MDA (green), read abundance of MALBAC (black), read abundance of WGA4 (blue), and read abundance of the bulk-cell sample (red). Each bar represents the total reads in a 500-kb bin, and all 5,797 bins are plotted. (**b**) Box plot representing the normalized reads in 5,7 bins in log10 scale. (**c-d**) Correlation between the three single-cell amplification methods and the bulk-cell sample without GC-correction (**c**) and with GC correction (**d**). For each 631-kb bin, the normalized read abundance of one sample is the x-value and the normalized read abundance of another sample is the y-value. The combination of these two values will give one dot on the plot for a particular bin. There are 4,580 bins in total. The 4,580 dots are plotted to show the reproducibility of the three single-cell genome amplification methods relative to an unamplified sample. A narrow distribution of dots along the y = x (red line) indicates good correlation between a single-cell method and the bulk-cell sample. For each method, one particular cell is chosen for this plot. **Supplementary Fig. S4** illustrates the results for all of the other cells.

To quantitatively measure the bin-to-bin variation, a box-plot was prepared for each of the cells used in our studies (Fig. 5b). Box plots characterize a sample using the 25th (Q1), 50th, and 75th (Q3) percentiles and the interquartile range (IQR = Q3-Q1). It covers the central 50% of the data. Quartiles are insensitive to outliers and preserve information about the center and spread. Consequently, they are preferred over the mean and standard deviation for population distributions. Among the three single-cell amplification methods, the IQR of the WGA4 method is the smallest, indicating that WGA4 has the least read fluctuation among the bins and the best performance with respect to coverage uniformity. We also plotted the normalized single-cell reads of each bin against results from bulk cells (Fig. 5c). The WGA4 method gives the best correlation (correlation efficiency 0.56) with bulk-cell samples. Random fragmentation of genomic DNA to around 300 bp in the WGA4 method may be responsible for the evenness of the whole genome amplification. Because MALBAC has a GC-bias toward high GC content (Fig. 3), the correlation between MALBAC and bulk-cell samples is poor. Therefore, we applied a GC-correction locally weighted scatterplot smoothing (LOWESS) algorithm (Baslan et al. 2012) to correct the GC-biases in all three single-cell methods and recalculated the correlation (Fig. 5d and Supplementary Fig. S7a,b). The correlation between MALBAC and bulk-cell samples was significantly improved, from 0.36 to 0.53. In our hands, the WGA4 method presents superior genome coverage uniformity and correlation with the bulk-cell sample. When single-cell genomes are amplified by MALBAC, after GC-correction, high-quality data can also be obtained.

### Detecting copy number variation

It has long been thought that all neurons in a brain share the same genome. However, recent evidence suggests that individual neurons could have non-identical genomes because of aneuploidy, active retrotransposons, and other DNA content variations(Rehen et al. 2005; Singer et al. 2010; Westra et al. 2010; Evrony et al. 2012). We applied the algorithm from Navin *et al*. to call CNVs in neurons(Baslan et al. 2012). In contrast to using fixed intervals to calculate copy number, we used bins of variable length with uniform expected unique read counts(Baslan et al. 2012). We also used the LOWESS algorithm(Benjamini and Speed 2012), which corrects for GC content bias. In our study, the MDA, MALBAC, and bulk-cell samples were prepared from rat embryo 2. WGA4 cell samples were prepared from rat 1. As shown in Fig. 6a, sequencing results from both the bulk-cell sample and the single neurons amplified by the MALBAC method showed one X chromosome, indicating that rat embryo 2 is a male. Single neurons from rat embryo 1 were amplified by the WGA4 method, and they showed two X chromosomes. Therefore, embryo 1 is a female. These results suggest that both the WGA4 and MALBAC methods can easily reveal chromosome-level CNVs. However, when the data from MDA were analyzed, it was challenging to identify the copy number of the X chromosome, even after aggressive binning into large genomic regions. We next investigated the CNV detection limit for the WGA4 and MALBAC methods. We developed a computational algorithm to evaluate the CNV detection limit (**Supplementary Table S2 and Table S3**). We performed the analysis for bin sizes that ranged from 63 kb to 12.7 Mb (average variable bin size). At the 641-kb bin size, CNVs were detected in all four WGA4 samples and two of five MALBAC samples. When the average bin size is reduced to 252 kb, CNVs were detected in one of four WGA4 samples and none of MALBAC samples. Therefore, both MALBAC and WGA4 may be used to detect CNVs with a bin size around 641 kb. Fig. 6a shows an example of the CNV detection data quality. At the 641-kb bin size resolution, the copy number patterns of most autosomal chromosome regions (Fig. 6a) from both the MALBAC and bulk-cell samples are identical. The bulk-cell sample has 10 CNVs (**Supplementary Table S3**). Among the 10 CNVs detected from the bulk-cell sample, eight CNVs are also detected in the MALBAC sample. Another two CNVs that are detected in the MALBAC sample are not observed in the bulk-cell sample. The WGA4 sample also has 10 CNVs, and eight of them are also observed in the bulk-cell sample (**Supplementary Table S3**). However, it should be emphasized that the WGA4 and bulk-cell samples are from two different embryos; therefore, the CNVs could be due to genomic variations between the two embryos. These comparative studies indicate that single-cell sequencing can reveal not only the information obtained from bulk-cell samples but also other fine details at the single-cell level. A recent study suggested that individual neurons may have somatic mosaic CNVs, in particular, aneuploidy in human postmortem brain samples(McConnell et al. 2013). We examined such a possibility in rats using our single-neuron sequencing results. As shown in Fig. 6b, a neuron (labeled WGA4 cell 6) displays a 20.5-Mb subchromosomal deletion in chromosome 8, which is not detected in another neuron (labeled WGA4 cell 5) or the bulk-cell sample. This result suggests that rat neurons also have mosaic CNV, which is consistent with report in human studies(McConnell et al. 2013).

**Figure 6.**
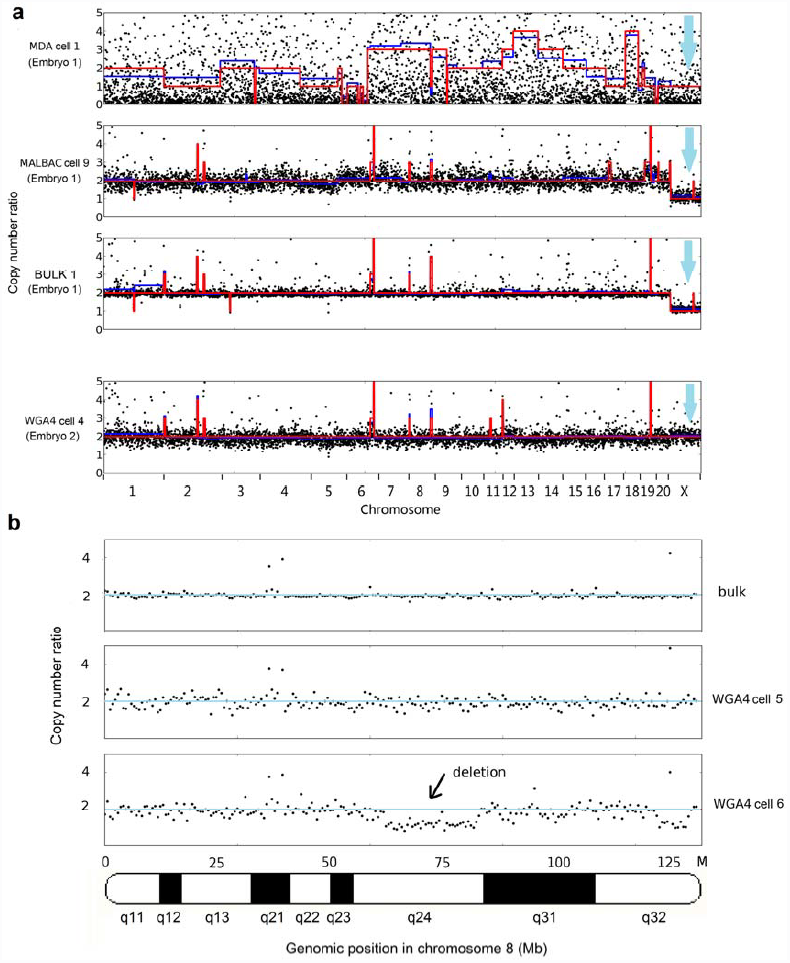
Copy number determination. (**a**) The copy numbers predicted for three single cells and the bulk-cell sample after GC-correction. The black dots are the normalized abundance of reads in 631-kb bins. The red line is the predicted copy number. MDA cell 1, MALBAC cell 9, and Bulk 1 are from rat embryo 2; WGA4 cell 6 is from rat embryo 1. The MALBAC and bulk-cell samples show one copy of the X chromosome, and the WGA4 sample shows two copies of the X chromosome. (**b**) Mosaic CNV is detected in rat neuron chromosome 8. The blue line is y = 2. The single-neuron WGA4 cell 6 has a subchromosomal deletion in chromosome 8 that is approximately 20.4 Mb in size.

## DISCUSSION

We quantitatively compared the performance of three single-cell whole genome amplification methods using rat hippocampal neurons as our study target. Using the bulk-cell sample as the benchmark, we have shown that the single-cell DNA sequencing varies in genome coverage, reproducibility, GC-bias, and coverage uniformity. MALBAC displays the best genome coverage with excellent reproducibility. WGA4 has the best performance in genome coverage uniformity. Both the WGA4 and MALBAC methods can detect the chromosome-level (Fig. 6a) and subchromosomal-level (Fig. 6b) CNVs.

Because MALBAC and WGA4 have their uniqueness and strengths, the information gained in this study will guide the selection of a proper single-cell genome amplification method, which depends on the chosen scientific questions to be addressed. In cancer biology, tumors display extensive somatic mutations and chromosome instability. Single-cell sequencing can be applied to assess the clonal structure of intra-tumoral heterogeneity. The MALBAC method has high genome coverage and may work better in detecting mutations than other methods(Zong et al. 2012). WGA4 has better genome coverage uniformity than other methods and can be applied to study chromosome instability and CNVs(Navin et al. 2011). In tumor metastasis, single-cell genome sequencing can be used to address the origin of metastasis and clinically monitor the metastasis by sequencing single circulating tumor cells(Ni et al. 2013). A combination of MALBAC and WGA4 can produce a comprehensive profile of genomic variations in tumors. In pre-implantation genetic diagnosis, we need to select embryos that have the greatest chance for a successful pregnancy and are free of monogenic disorders. MALBAC sequencing of a polar body enabled us to accurately detect aneuploidy and SNPs in disease-associated alleles(Hou et al. 2013). In neurobiology, neuronal diversity has been increasingly recognized to be mediated by somatic variations in the genome and epigenome, which mainly include chromosome instability, aneuploidy (rarely polyploidy), mosaic subchromosomal rearrangements, and intercellular changes in the epigenetic profile. Our results show that the WGA4 method can successfully detect aneuploidy and subchromosomal CNVs, which is consistent with the report from McConnell *et al.*(McConnell et al. 2013). In the field of microorganism genomics, many bacteria and archaea are difficult to culture, and single-cell sequencing is a powerful tool to profile their genomes. MALBAC will be at a unique position to sequence genomes with high GC content.

Although single-cell sequencing experimental techniques are rapidly developing, few bioinformatics tools are available that are specific to single-cell genomics analysis. Each single-cell amplification method has its own built-in pattern of biases, and future bioinformatics tools will need to consider these patterns(Ning et al. 2014). For example, MALBAC preferentially amplifies the high-GC-content regions, and this preference is highly reproducible. We can partially correct this bias by normalizing the coverage by the GC content. Better algorithms to identify CNVs are also needed, especially algorithms with criteria that define the resolution of the CNV detection limit of each method.

Single-cell sequencing techniques will be further improved in the future. For this, the quality of the single cell is vitally important. Here, we used micromanipulation with a micro glass pipette to select a single nucleus for amplification. A single nucleus is better for amplification than the whole cell because it contains fewer enzymes and proteins that may interfere with the amplification, thus reducing the amplification background. Additionally, reducing the reaction volume can improve the fidelity and reduce the bias of the method. Microfluidics(Wang et al. 2012) and nanoliter-based(Gole et al. 2013) single-cell amplification achieve better data quality than in-tube amplification. Also, single-molecule sequencing methods such as third-generation sequencing technologies eliminate the amplification step before sequencing(Harris et al. 2008; Eid et al. 2009). As such, they eliminate amplification bias and hold great promise for future single-cell sequencing studies.

## METHODS

### Primary hippocampal neuron culture and single-neuron nucleus isolation

As described previously(Zhang et al. 2009), hippocampi that were dissected from embryonic day 18 Sprague–Dawley rat embryos were digested with papain (0.5 mg/mL in Hank’s balanced salt solution HBSS, 37°C for 20 minutes), washed, and gently triturated by passing the tissue through a Pasteur pipette with a sterile tip. Neurons were counted and plated onto poly-L-lysine (Sigma, 0.5 mg/mL) pre-coated 60-mm Petri dishes (Becton Dickinson, Bedford, MA) at 2×10^6^ per dish to isolate DNA from a population of cells (2 million neurons) or dishes containing five glass coverslips (0.3×10^6^ per 60-mm dish) for single-neuron nucleus isolation. To ensure high-quality cell adhesion and growth, coverslips were first incubated in 100% nitric acid overnight, thoroughly washed with five changes of large amounts of deionized (DI) water, and stored in 70% ethanol. Coverslips were then flamed, dried, coated with poly-L-lysine (Sigma, 0.5 mg/mL) overnight, and washed three times with sterile DI water again before being incubated in plating medium for cell plating. The plating medium is 1× Minimum Essential Media (MEM, Cellgro)containing 10% fetal bovine serum, 5% horse serum (HS), 31 mg L-cysteine, and 1% pentamidine/streptomycin/gentamicin (P/S/G). Twenty-four hours after plating, the plating medium was replaced by feeding medium (Neurobasal medium from Cellgro supplemented with 1% HS, 2% Gibco B-27, and 1% P/S/G). Thereafter, neurons were fed twice per week with 2mL feeding medium per dish for 2 weeks until use. Given the important roles played by glial cells in neuron development and synaptogenesis, glial cell growth was suppressed by supplementing feeding medium with 5-flouro-2-deoxyuridine beginning on day *in vitro* (DIV) 5, but they were not completely eliminated from the culture.

A single neuron nucleus was extracted directly through micromanipulation using a micro glass pipette on an electrophysiological recording system. The micro glass pipettes were made on a flaming micropipette puller (Model P-97, Sutter Instrument) by pulling capillary glass tubing (Model G85150T-3, Warner Instruments). The flaming temperature and pulling velocity were adjusted accordingly to generate micro pipettes with tip diameters ranging between 5 and 10 μm. The micro pipette was then filled with 1× artificial cerebrospinal fluid and installed on the electrophysiological recording system. A micromanipulator (Model MP-225, Sutter Instrument) was employed to control the micro pipette to slowly approach the target neurons. Typical hippocampal pyramidal neurons were identified under a 32× objective (numerical aperture, 0.4) with a Zeiss Axiovert 100 microscope. Once in touch with the cell membrane, negative pressure was applied to gently inhale the whole nucleus from the neuron and into the micro glass pipette. The isolated cell (nucleus) in the micro glass pipette was injected into a 200-µL PCR-ready vessel with 3 μL Phosphate buffered saline (PBS, Sigma-Aldrich, Cat no. P5368-10PAK) which was free of DNase, RNase, and pyrogens.

#### Whole genome amplification

WGA4 amplification was performed on a single neuron nucleus as described in the Sigma-Aldrich GenomePlex WGA4 kit (Sigma-Aldrich, Cat no. WGA4-10RXN). Briefly, we first lysed the nucleus and removed the proteins by incubating the mixture at 50°C for 1 hour. The genomic DNA was fragmented for 4 minutes at 99°C. A set of random primers linked with common adaptors was annealed to the fragmented DNA template at the following series of temperatures: 16°C for 20 minutes, 24°C for 20 minutes, 37°C for 20 minutes, 75°C for 5 minutes, and 4°C hold. Then, PCR was performed to amplify the library with an initial denaturation at 95°C for 3 minutes, and 25 cycles of 94°C for 30 seconds and 65°C for 5 minutes. The PCR product was purified using the Qiagen PCR Purification kit. Most DNA in the library is from 200 bp to 400 bp. The MDA amplification was performed with the Qiagen REPLI-g Single Cell kit. We strictly followed the manufacturer instructions. The single-cell amplification by the MALBAC method was performed by Yikong Genomics (http://www.yikongenomics.cn/).

#### Sequencing library preparation and sequencing

After single-cell genomic DNA was amplified, the sequencing libraries were constructed by BGI-Shenzhen sequenced using the Illumina HiSeq 2000 sequencing platform.

#### Bioinformatics analysis

##### (A) Read alignment

The total number reads for each sample ranged from 8 million to 58 million. Because the MALBAC and WGA4 methods added around 30-bp adaptors to each read, we deleted the nucleotide sequences of adaptors and truncated the reads to 60 bp. This truncation was performed for all samples to ensure that all single-cell sequencing data were evaluated using reads of the same length. After filtering for clean reads, the data were mapped to the rat reference genome (rnt5) using Bowtie2 software with the default parameters. Duplicates were removed using SAMtools (Li et al. 2009) and MarkDuplicates from the Picard software suite.

##### (B) Genome coverage efficiency

For each sample, we randomly selected 10% of the reads from the total sequence reads. These reads were aligned to the reference genome. We calculated the percentage of the genome that was covered by at least one read. We repeated the sampling 10 times and determined the average genome coverage at 10% sampling depth. We did this by selecting 20%, 30%, …, and 90% reads. At different sampling depths, the same process was followed to generate the information shown in Fig. 2.

##### (C) GC-bias

To calculate the GC-composition of the reference genome, we divided the reference rat genome into continuous 60-bp windows. The GC content of each window was calculated. The frequency of reads of 1% of the GC-content intervals was counted. To calculate the GC-composition of single-cell samples and the bulk-cell sample, we determined the GC content of each sequencing read, which was used to calculate the relationship between the read distribution frequency and GC content. We normalized the total reads of each sample to 10 million.

Relative coverage is defined as the ratio of the normalized read number of a particular sample to the normalized read number of the reference genome. A relative coverage of 1 indicates that a particular base is covered at the expected average rate. A relative coverage above 1 indicates higher than expected coverage, and a relative coverage below 1 indicates lower than expected coverage. This was used to generate Fig 3b–d.

##### (D) Reproducibility

We divided the rat genome into 500-kb bins. The total bin number is 5,797. We then calculated the ratio between the number of reads of each bin and the average number of reads of all 5,797 bins. To quantitatively evaluate the reproducibility, a plot was generated using the ratio of each of the 5,797 bins from the first single cell as the x-value and the ratio of the same bin from the second single cell as the y-value. The correlation coefficient between the ratios of two single cells was calculated. For perfectly reproducible data, data points should all fall onto the y = x line. Hierarchical clustering was performed using the hcluster command in R language.

##### (E) LOWESS model of GC-correction

We employed the LOWESS model to perform GC-correction. First, the GC content and read count was calculated for each bin. Then, a local linear polynomial fit was performed for the GC content and read count. Finally, the regression value of the read count was used to replace the original read count for each bin. The LOWESS model for GC-correction has been systematically studied in previous reports (Baslan et al. 2012; Benjamini and Speed 2012). We adopted a LOWESS function in the R package for this study.

##### (F) Determine the boundary of bins with variable bin sizes

From the rat reference genome (rnt5), we simulated 60-bp sequence reads by sampling the genome at 60×. These sequences were mapped back to the rat reference genome using Bowtie2 with the default parameters. The chromosomes were then divided into bins with an equal number of simulated reads.

##### (G) Copy number calculation

We used the method proposed by Navin *et al.* to calculate CNV. Copy number was assessed with bins of variable size (641 kb on average). We first calculated the number of reads that were mapped to each bin. Then, we performed the GC-correction using the LOWESS model. We used the circular binary segmentation algorithm from an R package (DNA copy) to group adjacent bins into segments(Venkatraman and Olshen 2007). The copy number of each segment was calculated as the mean read number of bins in the segment divided by the mean read number of bins of all autosomal chromosomes (multiplied by 2). The copy number is shown as the blue line in Fig. 6a. We rounded the copy number to integers. The rounded copy number is shown as the red line in Fig. 6a.

## ACKNOWLEDGMENTS

The authors would like to acknowledge BGI-Shenzhen and Yikong Genomics Co. Ltd. for providing sequencing services. This work was supported by the National Natural Science Foundation of China (J.H., Grant No. 31200688) and in part by the National Science Foundation (P.L., CHE–1309148) and the National Institutes of Health (H.Y.M., MH079407).

